# Brain Oscillations during Visuomotor Processing in Children with ASD

**DOI:** 10.1101/2023.01.17.524474

**Authors:** Emily May Wilson, Kyung-min An

**Affiliations:** Centre for Human Brain Health, School of Psychology, University of Birmingham, Birmingham, B15 2TT, United Kingdom

## Abstract

Autism Spectrum disorder (ASD) has been reported to often involve significant motor dysfunction in addition to other core traits. While individual differences in visual processing have also been noted in individuals with ASD. However, the brain oscillations between visual and motor processing in ASD are poorly understood due to a lack of research surrounding the topic. To elucidate this, we focused on the power-power correlation between visual- and motor-related oscillations. We recorded magnetoencephalography in 18 children with ASD and 19 IQ-matched typically developing children aged 4-7 while they pressed a button during a visual-targeted motor task. We estimated visual- and motor-related brain oscillations and focused on the gamma oscillations. We observed gamma oscillatory power changes during the visual-targeted motor task in both groups. We further found statistically significant differences in motor-related gamma power in the right primary motor cortex, but not in the left primary motor cortex between the two groups. We finally found a significant correlation between the left visual gamma power (50-60 Hz, 150-450ms) and the left motor gamma power (70-90 Hz, 0-100 ms) in the TD group. The current findings revealed differences in visuomotor processing in individuals with ASD compared to TD children. These results may be helpful in understanding the neural mechanisms underlying visuomotor processing in ASD, supporting the development of a potential biomarker for ASD.

## Introduction

Autism Spectrum disorder (ASD) is a neurodevelopmental disorder characterized by difficulties in social interactions and communication, repetitive behaviors, and restricted interests (American Psychiatric Association, 2013). Furthermore, it has been previously reported that children and adults with ASD also present significant motor dysfunction with their main traits (Fournier et al., 2010; Lloyd et al., 2013; Lord et al., 2000; Teitelbaum et al., 1998). Motor abnormalities in ASD were first included in a definition in 1943 (Kanner, 1943) and have continued to be widely reported, including fine and gross motor skills, gait, balance and posture, which are all key for motor function in humans (Bryson et al., 2007; Fournier et al., 2010; Jansiewicz et al., 2006; Lloyd et al., 2013; Lord et al., 2000; Molloy et al., 2003; Noterdaeme et al., 2002; Radonovich et al., 2013). Previous studies have reported that 80% of children with ASD have overall motor skills impairment including gross and fine motor performance (Bryson et al., 2007; Fournier et al., 2010; Jansiewicz et al., 2006; Lloyd et al., 2013; Lord et al., 2000; Molloy et al., 2003; Noterdaeme et al., 2002; Radonovich et al., 2013). Motor dysfunction is not included as a key symptom in the diagnostic tools used for an official diagnosis, a plethora of studies have suggested that motor dysfunction should be considered a core symptom of ASD (Fournier et al., 2010; Lloyd et al., 2013). Brain oscillations are patterns of neural rhythmic activity, and motor control is linked to brain oscillations in the beta and gamma frequency bands (An et al., 2021; An et al., 2018).

While a wide range of studies focuses on the link between ASD and motor dysfunction, limited literature focuses on the connection between visual processing and ASD. It is currently suggested that individual differences exist in visual perception in individuals with ASD (Robertson & Baron-Cohen, 2017; Ronconi et al., 2020; Seymour et al., 2019). Several studies suggest that individuals with ASD excel at visual search and crowding tasks (Lindor et al., 2018; O’riordan et al., 2001; Ronconi et al., 2020). Individuals with ASD further have significantly larger cortical responses to stimuli in both the frontal and parieto-occipital areas of the brain during the early stages of visual processing (Baruth et al., 2010). Sun et al (2012) explored impaired gamma band activity during perceptual organization in adults with ASD. They used magnetoencephalography (MEG) and the Mooney face paradigm to explore if gamma band activity is linked to atypical brain functioning in ASD in adults. They analyzed the MEG data in the 25 to 150 Hz range and tried to identify sources of spectral power using a beamforming approach (Sun et al., 2012). They found that participants with ASD presented elevated reaction times and reduced detection rates during the perception of the Mooney face paradigm but the responses to the inverted stimuli were in the normal range. They further found that perceptual organization was linked to a reduction in both the amplitude and phase locking of gamma band activity in the ASD group. In addition, they found that in the control group, the perception organization of Mooney faces involved increased 60-120 Hz activity which was found in the frontoparietal network. However, stronger neural activation was found in the visual frontal (Sun et al., 2012). They suggest their findings show the contribution impaired gamma-band activity has on complex visual processing in ASD. With them further suggesting that individual with ASD present atypical modulation of high frequency (Sun et al., 2012). However, this study only focuses on high gamma frequency bands meaning exploration of lower frequency gamma bands needs to be explored.

However, a recent study conducted by Seymour et al., (2019) explored dysregulated oscillatory connectivity in the visual system in autistic adolescents. They used magnetoencephalography (MEG) and a visual grating paradigm to explore connectivity in the visual domain with ASD and TD adolescents. They found that both TD and ASD groups showed similar increases in gamma (40-80 Hz) frequency power and decreased alpha (8-13 Hz) frequency power in the occipital cortex (Seymour et al., 2019). However, Seymour et al., (2019) found differences between the groups after conducting a more detailed analysis of the data. They found that feedback between two areas of interest in the visual cortex (V4 to V1), which is mediated by alpha oscillations, is significantly reduced in the ASD group compared to the TD group. This reduction was further positively correlated to the autistic quotient scores, which were collected prior to the experiment. This is consistent with the atypical visual hierarchy, which is linked to ASD, which is seen to be defined by a reduction in top-down modulation of visual input by alpha-band oscillations (Seymour et al., 2019). In addition, in the V1, they found that the coupling of the alpha-phase to gamma amplitude was reduced in the ASD group compared to the TD group. These results suggest that visual processes in ASD are less adapted to contextual feedback information (Seymour et al., 2019). However, the study only focuses on two areas of interest, leaving the connections in the wider visual cortex unknown.

Following this, Ronconi et al., (2020) explored whether detail-orientated visual perception is linked with altered neural oscillations in both beta and alpha frequency bands. Children with a diagnosis of ASD and TD performed a visual crowding task. Through comparisons between the ASD and TD groups, they found decreased activity in the occipital, inferior-temporal and inferior/middle frontal regions in the ASD group during conditions needing detailed-orientated perception (Ronconi et al., 2020). Ronconi et al. (2020) further found that detail-oriented perception was linked to an event-related beta power reduction (15-30 Hz) in the TD group, which was not found in the ASD group. These results suggest that altered communication in beta frequency bands may clarify atypical perception in ASD. However, visual beta and gamma oscillations have not been widely linked to being associated with ASD. So, in this current study, we investigated visual- and motor-related brain oscillations using child-customized MEG, while child participants conducted visual-motor tasks.

Recently there has been increasing interest in the connection between visual and motor processing in individuals with ASD. However, there is limited literature surrounding visual-motor processing in individuals with ASD. A previous study using resting state fMRI (functional magnetic resonance imaging) has suggested that there is intrinsic asynchrony regarding visual and motor systems in children with ASD (Nebel et al., 2016). Children with highly out-of-sync intrinsic visual-motor connectivity presented more severe ASD traits. This is suggested to contribute to impaired social and communication skills, which are noted as key characteristics of ASD. However, it is still unclear what the trajectory is of visual-motor development in children with ASD (Nebel et al., 2016). In addition, it has been suggested that visual-motor integration (VMI) is impaired with individuals with ASD being biased towards somatosensation (Proprioceptive feedback) compared to visual feedback (Lidstone & Mostofsky, 2021). Lidstone and Mostofsky (2021) suggest that people with ASD struggle to incorporate certain amounts of visual information to help update predictive motor control representations. They found that the brain regions implicit in prediction and updating representations such as the cerebellum and the Inferior parietal lobule show atypical connectivity and structures in individuals with ASD (Lidstone & Mostofsky, 2021). They conclude that motor imitation is impaired in individuals with ASD. With future studies being crucial in understanding the key mechanisms which underlie VMI in individuals with ASD, with the goal of creating targeted therapeutic interventions to support those with ASD (Lidstone & Mostofsky, 2021). However, the lack of literature exploring this connection with neurophysiological techniques which measure brain oscillations, such as MEG and OPM (Optically pumped magnetometers), has left vast gaps in fully understanding visual-motor processing in individuals with ASD.

In the present study, we measured visual- and motor-related brain oscillations using child-customized MEG, while child participants conducted visual-motor task. To investigate the brain, we focused on the power-power correlation between visual- and motor-related oscillations. We hypothesized that visual and motor oscillatory correlation would be altered in children with ASD.

## 2. Materials and methods

### 2.1. Participants

Eighteen children with ASD (mean = 6.00 years, SD= 0.59, 5 females, 13 males) and 19 typically developing children who were matched using age participated in this study (Mean - 5.71, SD=0.46, four females and 15 males). Previous data from An et al. (2021) was used for the current study. All participants were identified as right-handed using the Edinburgh Handedness Inventory (Oldfield, 1971). Participants in the ASD condition had a confirmed diagnosis of autism spectrum disorder based on the criteria for either the DSM-V for autism and Asperger’s syndrome (American Psychiatric Association, 2013), the Diagnostic Interview for Social and Communication Disorders (Wing et al., 2002) and/or the Autism Diagnostic Observational Schedule (ADOS) (Lord et al., 2000). In addition to this, the Kaufman Assessment Battery for Children (K-ABC) was used to assess the intelligence of the participants. This presented no significant differences between the two groups when it came to achievement scores of the K-ABC (*t(35)= 1.439, p =0.159, Cohen’s d = 0.473*). All diagnoses were conducted by licensed psychiatrists and clinical speech therapists. The Kanazawa University Hospital Ethics committee approved this study and written informed consent was gained from all the parents of the participants.

### 2.2. Experimental Paradigm

To investigate the visual- and motor-related oscillations, we used a video-game-like motor task developed and used by our previous studies (An et al., 2018; An et al., 2021. The game-like motor task was developed using Presentation software (Neurobehavioral systems). The video-game-like motor task includes 100 visual targets and 100 button presses. The main aim of this visual-motor task is to help a puppy collect fruit by pressing a button. After 1200 ms, the puppy ran to the left side of the screen with the fixation dot being located in the middle of the screen. The participants were instructed to gaze at a fixation point which was presented in the middle of the screen to help reduce artefacts due to eye movement. The participants were then instructed to press the button with their right index finger when the fruit arrived at the fixation point in the middle of the screen as the visual target. When the participant pressed the button, the puppy jumped to the fruit and caught the fruit. The fruit targets randomly appeared every 3.5 to 4.5 seconds after the participants pressed the button. There were ten trials in each block, with ten blocks in total. As a reward for successfully completing each block, a bone with a red ribbon was presented to the puppy to help encourage the participants.

The motor task we designed was created to help minimize the participant’s eye movements by displaying a fixation point in the middle of the screen. We instructed participants to press the button as quickly as possible, this allowed us to calculate the response time. A non-magnetic fibre optic response pad (LUMINA LU400-PAIR, Cedrus Corporation, San Pedro, CA, USA) was used to measure participants’ button responses. Before beginning the motor task, the participants were instructed to hold the button response pad and r est their right Index finger on the response button.

During the motor task, MEG was recorded for approximately 9 minutes (100 button presses). The visual stimuli were presented on a screen (26◦ × 21◦ of visual angle) using an LCD projector (IPSiO PJWX6170N, Ricoh Company, Ltd., Tokyo, Japan). The visual stimuli were presented at random every 3.5-4.5 seconds after the button press response. If participants failed to detect the visual target, the puppy would fall down, and the trail was then repeated. The failed trials were not included or used for data analysis.

### 2.3. Magnetoencephalography recording

A whole-head 151-channel child customized MEG system was used to acquire MEG data (PQ 1151 R, Yokogawa/KIT, Kanazawa Japan) in a magnetically shielded room. The MEG signals were digitized at 2000 Hz and low-pass filtered at 200 Hz. Prior to the experiment, we explained the experimental paradigm and procedures to the participants and their legal guardians. The participants were shown one practice block of the motor task to help familiarize themselves with the paradigm and experiment. Participants were recorded during MEG measurement in a comfortable supine position on a bed with two experimenters at the bedside to support and maintain the participant’s attention on the task. Four head positioning coils were placed on the participant’s scalp (Cz 5 cm anterior from Cz and 5 cm from the superior side of the left and right pre-anterior points). The location of the positioning coils was measured before the participant’s MEG recordings to calculate the participant’s head position relative to the MEG sensors. A video monitoring system was used to monitor the participants and assess the participant’s compliance with our instructions. It was also used to help record any noticeable artefacts such as inappropriate head position, head motion or inconsistent attention to the screen. A 3D digitizer (Fastrak Polhemus, Colchester, VT, USA) was used to measure inconsistent screen attention, head shape and the location of the positioning coils. An MRI scan was conducted using T1-weighted gradient echo and Silenz pulse sequence (repetition time [TR] = 435.68 ms, echo time [TE] =- o.024 ms, flip angle – 7 degrees, field of view [FOV] = 220mm, matrix size = 256 x256 pixels, slice thickness – 1.7 mm and 130 transaxial images in a 1.5-tesla Magnetic resonance imaging scanner (SIGNA Explorer, GE Healthcare, USA).

### 2.4. Data analysis

The child MEG data were processed using the MATLAB (Mathworks, Natick, MA, USA) and Brainstorm toolbox (Tadel et al., 2011). The continuous MEG data were band-pass filtered from 0.3 to 200 Hz and notch filtered at 60, 120, and 180 Hz. We applied an independent component analysis method (“RunICA” implemented in Brainstorm, www.sccn.ucsd.edu/eeglab/) to remove artifacts caused by eye blinks, eye movements, and heartbeats. We identified the independent components representing cardiac and ocular activity by visual inspection based on time course and topography. We rejected the components representing artifacts and the remaining components were back projected into the signal space. We segmented the artifact-rejected data from –3 to 3 s relative to the visual target onset and button-press onset. We automatically rejected trials exceeding 2000 fT/cm for MEG gradiometer channels and removed trials with small muscle artifacts by visual inspection.

For the source localization, we estimated the weighted minimum norm estimates (wMNE), which is implemented in the Brainstorm toolbox (Hämäläinen & Ilmoniemi, 1994; Hauk, 2004; Lin et al., 2006). An overlapping sphere head model was calculated using individual MRI images, and the noise-covariance matrix were estimated for each participant during the baseline period (−2 to –1.5 s according to the button-press onset and –0.6 to –0.1 s according to the visual target onset). We calculated wMNE source localization using the overlapping-sphere head model with a Tikhonov regularization (λ = 0.1) (Tikhonov & Arsenin, 1977). We projected the individual cortical sources into the ICBM152 template anatomy in MNE coordinates.

We applied a bandpass filter to all preprocessed data between 0.3 to 30 Hz and averaged trials for each participant to calculate motor fields (MFs) and visual-evoked fields (VEFs). We calculated motor-fields (MFs) by averaging the trials segmented by the button press onset of all participants and further normalizing by the baseline period of –2 to –1.5 seconds. Grand-averaged MFs were calculated by averaging individual MFs across all participants and the peak of the MFs was shown at 20 to 40 ms according to the button response. The cortical sources of individual MFs were calculated using wMNE and were projected on the ICBM152 template anatomy in MNI coordinates. We could see that cortical sources of the MFs were shown in the bilateral primary motor cortex (M1). We calculated VEFs by averaging the trials segmented by visual target onset of all participants and further normalizing the baseline period of –0.6 to –0.1 seconds. Grand-averaged VEFs were calculated by averaging individual VEFs across all participants and the peak of the VEFs was shown at 260 to 280 ms according to the visual target. The cortical sources of the individual VEFs were calculated using wMNE and were projected on the ICBM152 template anatomy in MNI coordinates. We found that the cortical sources of the VEFs were shown in the bilateral cuneus. For further analysis, we selected the bilateral cuneus and M1 from the Desikan-Killiany atlas (Desikan et al., 2006), which is defined using FreeSurfer version 6.0 (http://surfer.nmr.mgh.harvard.edu/).

To determine the visual- and motor-related oscillatory changes, time-frequency representations (TFRs) were calculated at 1 to 100 Hz using a seven-cycle Morlet wavelet for each single trial source data. We calculated the percentage change in power relative to the baseline by converting the TFRs. The TFRs of the trial source was averaged within participants and also group averaged across both ASD and TD participants. We observed power increases in motor gamma oscillations from 70 to 90 Hz at 0 to 100ms according to button response onset. We found power increases in visual gamma oscillations from 50 t o 60 Hz at 150 to 450 ms according to visual target onset. The power change was then calculated by averaging the values in the time-frequency windows and we defined power of motor gamma and visual gamma oscillations.

### 2.5. Statistical analysis

We used SPSS VERSION 28.0 (IBM Corporation, New York) for our statistical analysis. We applied independent t-tests to compare differences between ASD and TD participants in behavior, motor, and visual conditions. To test the hypothesis, we completed comparisons conducted with both TD and ASD groups, TD separately and ASD separately. Comparisons were conducted with both ASD and TD groups, comparing participant characteristics, behavior and motor, behavior and visual, and visual and motor conditions. Next, comparisons were conducted on only the TD group, comparing participant characteristics, visual and motor, behavior and visual and motor and behavioral conditions. Finally, comparisons were conducted using only the ASD group, comparing participant characteristics, visual and motor, visual and behavioral and motor and behavioral conditions. Scatter graphs were then created to help establish positive correlations on significant results.

## 3. Results

### 3.1. Evoked brain response during visual-motor task

We calculated individual VEFs and MFs and grand-averaged across all participants. Figure 1A demonstrates the grand-averaged VEFs across all participants. The maximum peak of the grand-averaged VEFs was found at 260 to 280 ms. At the maximum peak of the VEFs, the field distributions in the MEG sensor space showed a dipole pattern on the occipital area (Figure 1B). The cortical sources at the peak of the VEFs were found in the bilateral cuneus (Figure 1C).

**Figure 1.**
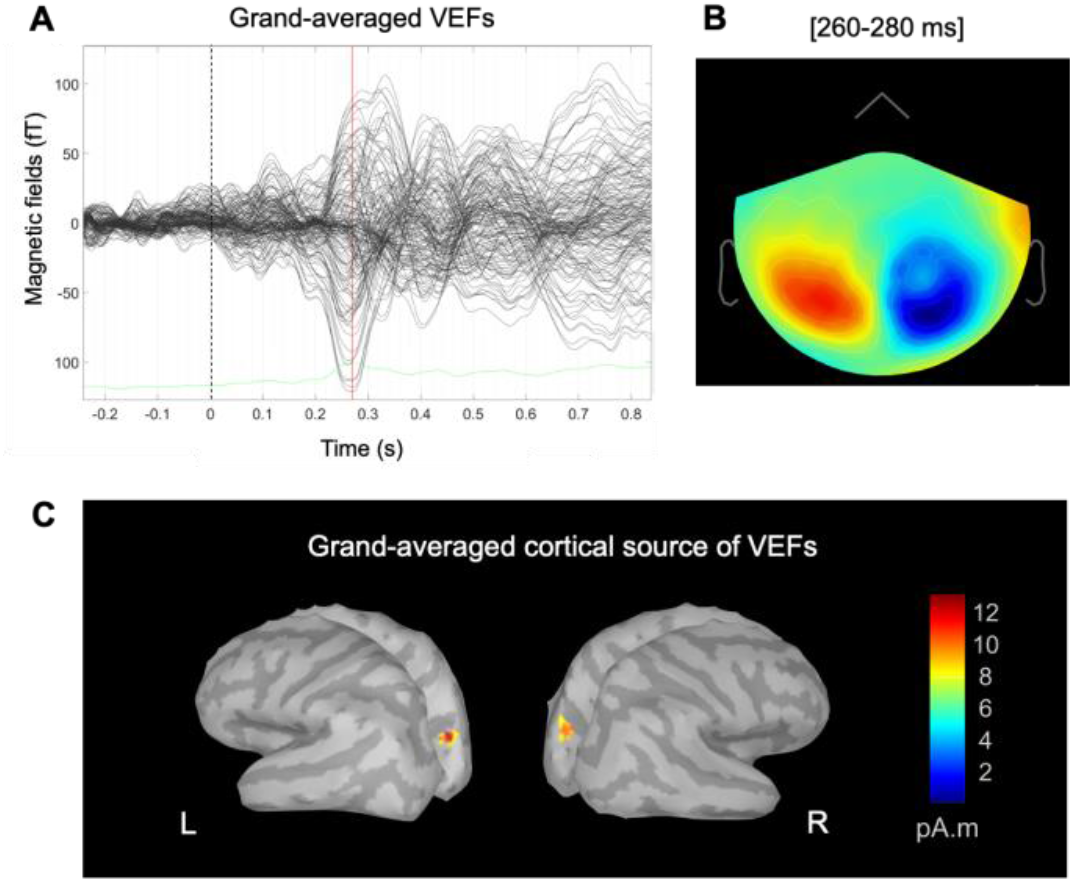
Grand averaged visual-evoked fields (VEFs), field distribution of VEFs in the MEG sensor space at the peak of the VEFs (260 to 280 ms), and its cortical sources.

Figure 2A demonstrates the grand-averaged MFs across all participants. The maximum peak of the grand-averaged MFs was observed at 20 to 40 ms. At the maximum peak of the MFs, the field distributions in the MEG sensor space showed a bilateral magnetic field pattern in the central area, which is shown left-lateralized (Figure 2B). The cortical sources at the peak of the MFs were found in the bilateral cuneus (Figure 2C).

**Figure 2.**
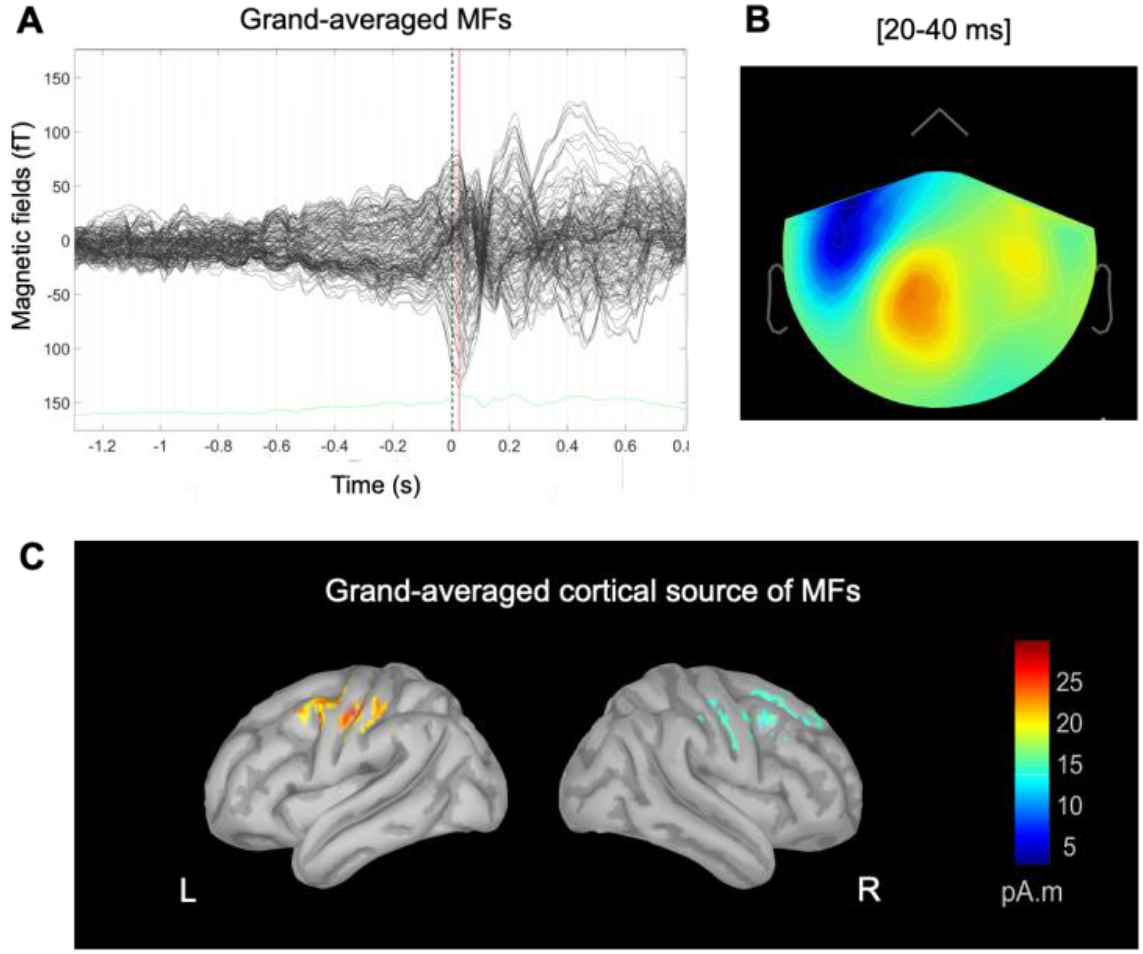
Grand averaged motor fields (MFs), field distribution of MFs in the MEG sensor space at the peak of the MFs (20 to 40 ms), and its cortical sources.

### 3.2. Bain oscillations during visual-motor task

We observed oscillatory power changes in TD children and children with ASD during the visual-motor task. Figure. 3 shows group-averaged time-frequency representations from the bilateral cuneus during visual stimuli in the TD and ASD groups. We found the power increase in visual gamma oscillations, which peaked at 50 to 60 Hz at 150 to 450 ms according to the onset of the visual target.

**Figure 3.**
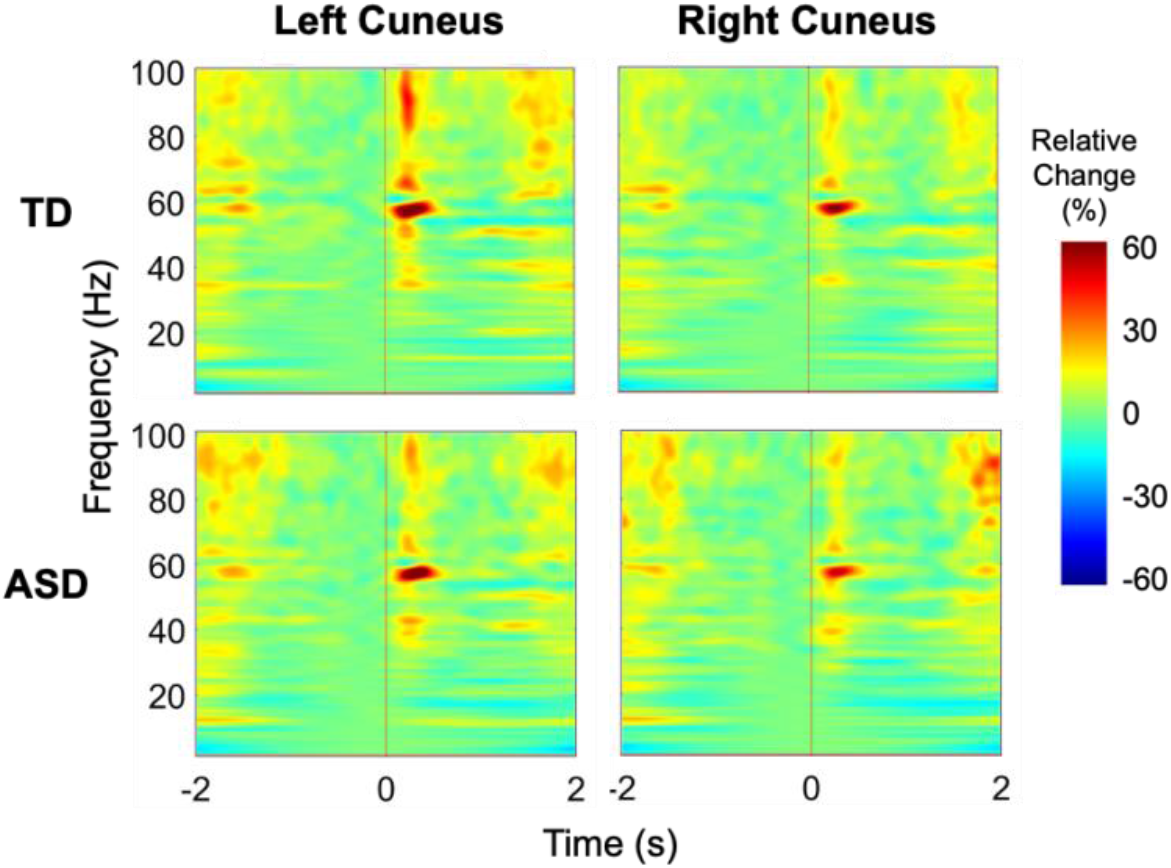
Grand-averaged time-frequency plots induced by visual stimuli in the bilateral cuneus from the TD and ASD groups.

Figure. 4 shows group-averaged time-frequency representations from the bilateral M1 during button response in the TD and ASD groups. As our previous studies (An et al., 2018; An et al., 2021), we found the power increase in movement-related gamma oscillations, which peaked at 70 to 90 Hz and 0 to 100 ms according to the onset of the button response in all participants.

**Figure 4.**
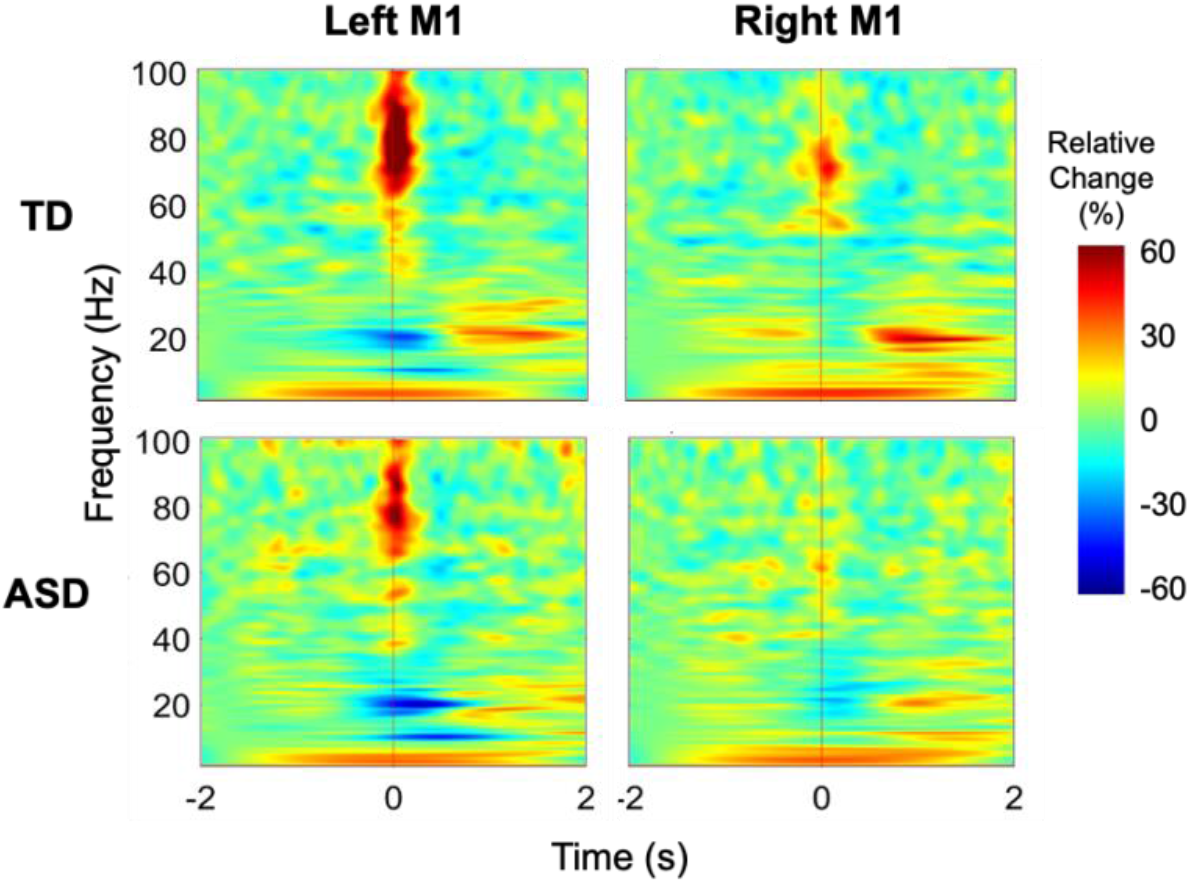
Grand-averaged time-frequency plots induced by button press in the bilateral primary motor cortex from the TD and ASD groups.

As same with the previous paper, there were statistically significant differences i n motor-related gamma power in the right primary motor cortex (*t(35) = 2.412, p = 0.021*), but not in the left primary motor cortex (*t(35) = 1.442, p = 0.158*) between the two groups (Figure 5A). However, we could not find any significant differences in the visual gamma power between the two groups in the bilateral cuneus (*t(35) = 0.570, p = 0.572* for the left cuneus, *t(35) = -0.307, p = 0.761* for the right cuneus) (Figure 5B).

**Figure 5.**
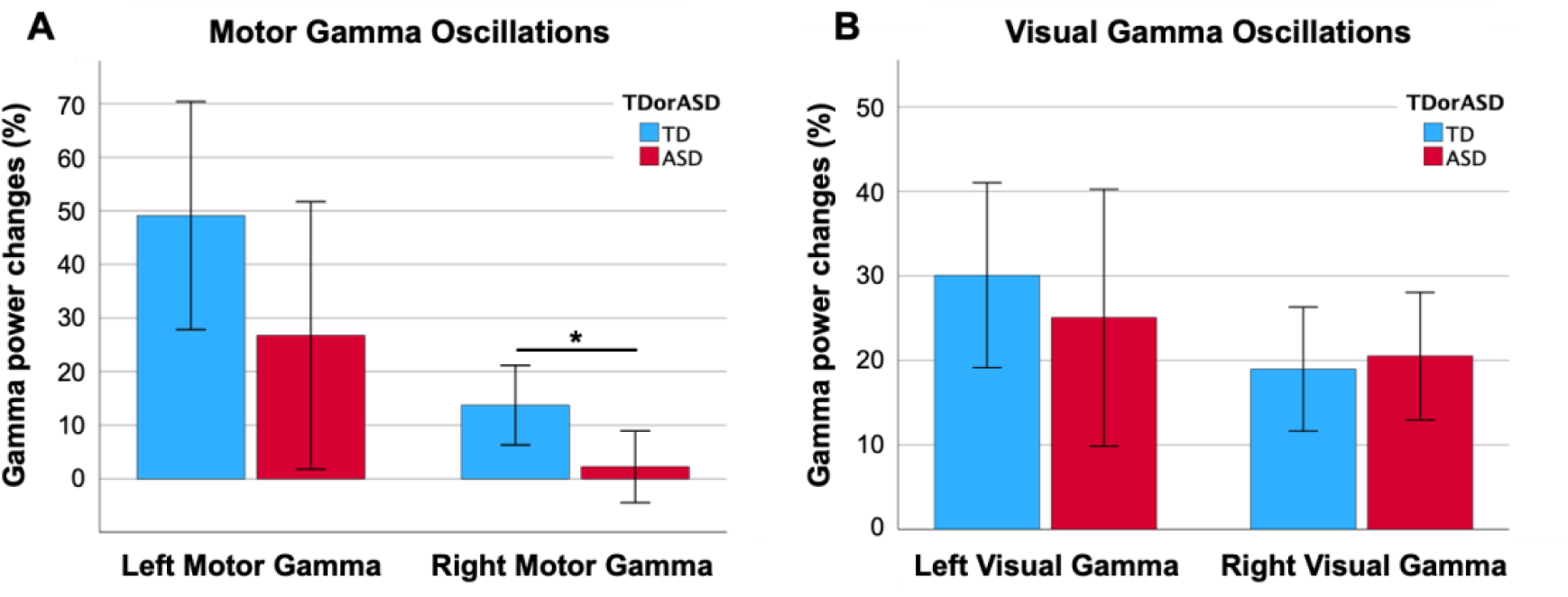
Comparison of the bilateral gamma power changes induced by button response and visual stimuli in TD and ASD groups respectively. The power of the motor-related gamma oscillations was calculated by averaging those in the time-frequency window at 70 to 90 Hz and 0 to 100ms. The ASD group presented smaller motor gamma power increases compared to the TD group in the right M1 (*t(35) = 2.412, p = 0.021)*, but not in the left M1 (*p>0.05*). The power of the visual gamma oscillations was calculated by averaging those in the time-frequency window at 50 to 60 Hz and 150 to 450ms. There were no significant differences in the bilateral visual gamma power between the two groups (*p>0.05*). The error bars represent a 95% confidence interval for each bar.

### 3.3. Correlations between visual- and motor-induced gamma power

We further calculated correlations using spearman’s rho correlations between the visual-induced gamma power and motor-induced gamma oscillations. We found a significant correlation between the left visual gamma power and the left motor gamma power in the TD group (ρ = -0.553, p = 0.014) (Figure 6), but not in the ASD group (ρ = 0.104, p = 0.681). There were no statistically significant correlations between visual and motor gamma power in the right hemisphere in the TD group (ρ = -0.388, p = 0.101) and in the ASD group (ρ = 0.011, p = 0.964).

**Figure 6.**
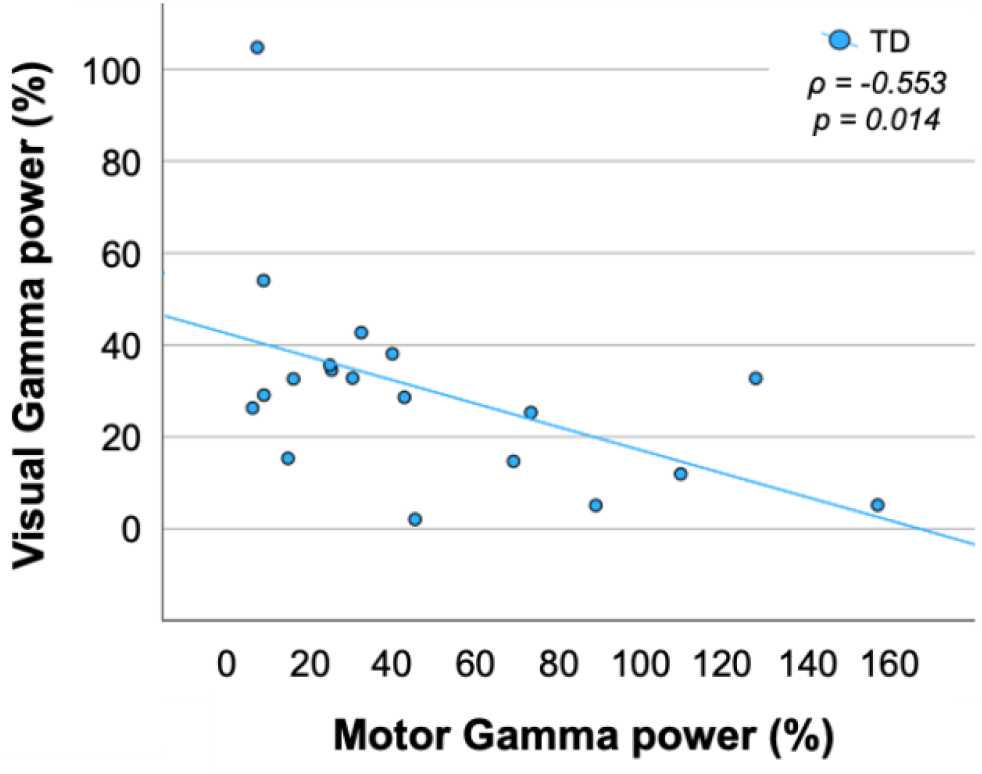
Scatter plot showing the correlation between the visual-induced gamma power (50 to 60 Hz, 150 to 450 ms) and Motor-related gamma power (70 to 90 Hz, 0 to 100ms). The negative correlation between the left visual-induced gamma power and motor-induced gamma power is observed (Spearman’s *ρ = -0.553, p = 0.014*).

## 4. Discussion

We investigated visual- and motor-related gamma oscillations and their relationship during visual-motor task to explore the brain mechanisms underlying visual-targeted motor processing in children with ASD. To our knowledge, this is the first neurophysiological study to focus on the brain oscillations related to visual-targeted motor processing in child participants. We measured power changes in the brain oscillations in the bilateral M1 and cuneus, while the child participant conducted the visually targeted motor tasks. During the task, we found both motor-induced and visual-induced gamma power increases in the TD and ASD groups. We confirmed that children with ASD showed decreased power in the motor-related gamma oscillations in the ipsilateral M1. However, we could not find any differences in the visual-induced gamma oscillations from the bilateral cuneus between the TD and ASD groups. In addition, we investigated the relationship between the motor gamma power and visual gamma power, and we found a significant correlation between the left visual gamma and left motor gamma in the TD group, but not in the ASD group.

### 4.1. Gamma oscillations related to visual stimuli

We found a power increase in gamma oscillations in the bilateral cuneus, which peaked at 50 to 60 Hz and 150 to 450 ms according to the onset of the visual target stimuli in the two groups. The visual gamma oscillations are known to play a critical role in temporal binding during sensory processing. Gamma band oscillations have been seen to be important for exploring deficits in temporal binding in individuals ASD (Seymour et al., 2019). Previous studies found that the power increase of the gamma frequency bands is related to the visual stimuli (Seymour et al., 2019; Sun et al., 2012). With elevated gamma activity in the visual regions which have been suggested to present the perception qualities of individuals with ASD (Sun et al., 2012).

We could not find any significant differences in the visual gamma power in the bilateral cuneus between the two groups. Our findings indicate that the ASD group does not exhibit alterations in the visual-related gamma increase in the bilateral cuneus. This finding conflicts with the previous MEG studies exploring visual oscillations in individuals with ASD. Previous studies have suggested that individuals with ASD show alterations in visual gamma power. Previous studies have found alterations in visual gamma power, Sun et al., (2012) found alterations in visual gamma power when it came to the perceptual organization in adults with ASD. They observed that visual perception was linked to significant impairments in gamma-band power in adults with ASD, suggesting an impairment in the timing of neural activity in adults with ASD (Sun et al., 2012). They further suggest that adults with ASD present significant impairments in the 60 to 120 Hz frequency range especially in the front posterior cortical networks. It is suggested that the 60 to 120 Hz gamma frequency range underlines perceptual processing in controls (TD participants), but this is significantly altered in adults with ASD (Sun et al., 2012). In addition, Seymour et al., (2019) found that individuals with ASD show alteration is visual gamma power. They found that both TD and ASD groups showed similar gamma frequency power increases (40 to 80 Hz) and decreased alpha frequency power (8 to 13 Hz) in the occipital cortex (Seymour et al., 2019). However, they further found that alpha-phase to gamma amplitude coupling was reduced in adolescents with ASD. They further suggest that the coupling between alpha and gamma oscillations in the visual cortex during perception is reduced in adolescents with ASD. They suggest that visual processing in ASD is less adapted to contextual feedback information (Seymour et al., 2019).

Previous literature has suggested that individuals with ASD have a significantly larger cortical responses to visual stimuli in the frontal and parieto-occipital and visual cortex of the brain (Baruth et al., 2010; Ronconi et al., 2020; Seymour et al., 2019; Sun et al., 2012) Like our findings. We found larger cortical responses in the bilateral cuneus which is located posterior the parieto-occipital and in the visual cortex of the brain.

Previous studies have shown increased visual oscillations when completing a visual task paradigm (Ronconi et al., 2020; Seymour et al., 2019; Sun et al., 2012). In addition, increases in visual gamma oscillations at 40-80Hz have been previously reported to be found in adolescents with ASD and those categorized as TD (Seymour et al., 2019). But it has been further reported that adults with ASD have impairments in the 60 to 120Hz range, especially in the frontoposterior cortical networks (Sun et al., 2012). While previous work that has explored visual oscillations in individuals with ASD has been mainly conducted on adolescents or adult participants, our results present different patterns but in young children. Previous studies have reported that adults and adolescents with ASD showed alterations in visual gamma (Seymour et al., 2019; Sun et al., 2012). We observed oscillatory gamma power changes in both TD and ASD groups. However, we could not find any significant differences in visual gamma power between the two groups in the bilateral cuneus. However, this could be explained by the age differences between participants in our study compared to previous literature exploring visual gamma oscillations in ASD. While previous studies have been conducted on adults (Seymour et al., 2019) and adolescents (Sun et al., 2012) with ASD, our study used participants who are children with ASD which may explain the differing results. This means our finding may be the result of using participants with a different age range compared to previous studies (Seymour et al., 2019; Sun et al., 2012). This suggests that children with ASD may present different alterations in visual gamma oscillations compared to adolescents and adults with ASD.

Individuals with ASD are seen to have an atypical visual hierarchy which is defined by top-down modulation by alpha-bands and also reduced coupling of alpha-phase to gamma amplitude (Robertson & Baron-Cohen, 2017; Seymour et al., 2019). However, in this study, we only focused on visual gamma oscillations and motor gamma and beta oscillations. But it is noted that detail-oriented perception is connected to event-related beta power reduction at 15-30 Hz in children who are classified as TD (Ronconi et al., 2020). Atypical perception in children with ASD is linked to altered communication in beta frequency bands. We observed motor-induced oscillations and visual-induced oscillations in young children with ASD and TD children.

### 4.2. Gamma oscillations related to movement

We further observed a power increase in gamma oscillations which peaked at 70-90 Hz at 0-100 ms according to the onset of the button response in all participants. However, the power of gamma oscillations is seen to increase directly after movement which has been previously reported in children (Cheyne & Ferrari, 2013) and adults (Cheyne et al., 2008; Cheyne & Ferrari, 2013; Muthukumaraswamy, 2010). This gamma increase is seen to be linked to the initiation of movement and motor execution (An et al., 2021; An et al., 2018). The results of this study supports our previous studies that indicates altered motor-related gamma activity is linked to impaired movement initiation in individuals with ASD.

We further found significant differences in motor-related gamma power in the right primary motor cortex but not in the left primary cortex. However, while motor dysfunction is excluded from diagnostic criteria for ASD currently, a plethora of studies have suggested that motor dysfunction should be classed as a core symptom of ASD (Fournier et al., 2010; Lloyd et al., 2013; London, 2014). A potential biomarker for ASD is suggested using gamma oscillations due to the significant differences in motor related gamma oscillations, with children with ASD presenting decreased motor-related gamma oscillations compared to TD children (An et al., 2018).

Our results support and replicate findings of our previous studies (An et al., 2021; An et al., 2018) which found reduced motor-related gamma oscillations in those with ASD. The findings further support the idea that motor dysfunction should be considered a key characteristic of ASD (An et al., 2021).

### 4.3. Oscillatory coupling during visual-targeted motor processing

We observed statistically significant correlations between the two groups when it came to motor gamma and visual gamma oscillations. We found significant correlations between the visual gamma oscillations which peaked at 50-60 Hz at 100-500 ms and motor gamma oscillations which peaked at 70-90 Hz at 0-100 ms. The results revealed significant correlations between left visual oscillations and left motor oscillations in children who were categorized as belonging to the TD group. However, this result was not replicated in the ASD group. A previous study highlighted that detail-orientated perception was linked to a beta power reduction at 15-30 Hz in children who are TD, but this was not seen to be replicated in children with ASD (Ronconi et al., 2020). Previous imaging studies have noted that children with highly out-of-sync intrinsic visual-motor connectivity present more sever ASD traits (Nebel et al., 2016). However, while significant correlations were only found in the TD group, it has been noted that in visual-motor integration (VMI) there is a bias towards proprioceptive feedback compared to visual feedback (Lidstone & Mostofsky, 2021). It has further been suggested that individuals with ASD have motor imitation impairment. This supports our findings due to us also finding impairments in motor-related gamma activity which is linked to impaired movement initiation in individuals with ASD. However, there are no studies which use neurophysiological techniques to explore visuomotor oscillations on both TD and ASD participants, with limited literature on this exploring visual- and motor-related brain oscillations in ASD.

## Conclusion

There are a number of limitations of our study which should be considered. First, due to the study’s only focusing on beta and gamma oscillations, the results do not take into account the role of other neural oscillations during visuomotor processing in ASD. Future studies should consider the role of additional neural oscillations on visuomotor processing. Secondly, our study contained a small sample of participants. This means the results are not sufficient for drawing firm conclusions around visuomotor processing in ASD. Future studies should consider recruiting a larger sample of participants with a larger age range to improve the reliability of findings.

Differences in motor and visual oscillations in ASD have previously only been observed separately. In the present study we demonstrated oscillatory power changes during the visual-motor task in both groups. In addition, the current study revealed significant differences in motor-related gamma power in the right primary motor cortex, but not in the left primary motor cortex between the two groups, which replicates the figs of previous studies. The findings extend the current understanding of motor dysfunction in individuals with ASD. However, our results further revealed a significant correlation between the left visual gamma power and the left motor gamma power in the TD group, but not the ASD group. The current findings revealed differences in visuomotor processing in individuals with ASD compared to TD children. The findings provide neurophysiological evidence for differences in visuomotor processing in children with ASD, supporting the development of a potential biomarker for ASD.

## Notes

### Competing Interest Statement

The authors have declared no competing interest.

